# The alarmone (p)ppGpp is part of the heat shock response of *Bacillus subtilis*

**DOI:** 10.1101/688689

**Authors:** Heinrich Schäfer, Bertrand Beckert, Wieland Steinchen, Aaron Nuss, Michael Beckstette, Ingo Hantke, Petra Sudzinová, Libor Krásný, Volkhard Kaever, Petra Dersch, Gert Bange, Daniel Wilson, Kürşad Turgay

## Abstract

Here, *B. subtilis* was used as a model organism to investigate how cells respond and adapt to proteotoxic stress conditions. Our experiments suggested that the stringent response, caused by raised levels of the (p)ppGpp alarmone, plays a role during thermotolerance development and the heat shock response. Accordingly, our experiments revealed a rapid increase of cellular (p)ppGpp levels upon heat shock as well as salt- and oxidative stress. Strains lacking (p)ppGpp exhibited increased stress sensitivity, while raised (p)ppGpp levels conferred increased stress tolerance to heat- and oxidative stress. During thermotolerance development, stress response genes were highly up-regulated together with a concurrent transcriptional down-regulation of the rRNA, which was influenced by the second messenger (p)ppGpp and the transcription factor Spx. Remarkably, we observed that (p)ppGpp appeared to control the cellular translational capacity and that during heat stress the raised cellular levels of the alarmone were able to curb the rate of protein synthesis. Furthermore, (p)ppGpp controls the heat-induced expression of Hpf and thus the formation of translationally inactive 100S disomes. These results indicate that *B. subtilis* cells respond to heat-mediated protein unfolding and aggregation, not only by raising the cellular repair capacity, but also by decreasing translation involving (p)ppGpp mediated stringent response to concurrently reduce the protein load for the cellular protein quality control system.

**Author Summary:** Here we demonstrate that the bacterial stringent response, which is known to slow down translation upon sensing nutrient starvation, is also intricately involved in the stress response of *B. subtilis* cells. The second messengers (p)ppGpp act as pleiotropic regulators during the adaptation to heat stress. (p)ppGpp slows down translation and is also involved in the transcriptional down-regulation of the translation machinery, together with the transcriptional stress regulator Spx. The stress-induced elevation of cellular (p)ppGpp levels confers increased stress tolerance and facilitates an improved protein homeostasis by reducing the load on the protein quality control system.

## Introduction

Bacteria have evolved complex and diverse regulatory networks to sense and respond to changes in the environment, which can include physical stresses or nutrient limitations [1]. The protein quality control system (PQS) comprises a conserved set of chaperones and proteases that monitor and maintain protein homeostasis is present in all cells. Various physical stresses, such as heat stress, favor the unfolding and aggregation of cellular proteins, which can be sensed by heat shock response systems, allowing an appropriate cellular stress response. This response includes the induction of the expression of chaperones and proteases of the PQS, also known as heat shock proteins [2, 3].

Interestingly, in all cells including *B. subtilis*, a short exposure to a raised but non-lethal temperature induces thermotolerance, an acquired resistance to otherwise lethal temperatures. Investigating the adaptation to such adverse conditions, also known as priming, allows the molecular mechanisms and interplay of the various cellular processes involved in the cellular stress and heat shock response to be studied [4, 5]. In *B. subtilis*, the heat shock response is orchestrated by multiple transcriptional regulators, including the heat-sensitive repressors HrcA & CtsR, which control the expression of the PQS and other general stress genes [6, 7]. HrcA regulates the expression of chaperones, while CtsR controls the expression of the AAA+ protease complexes [8–10]. The general stress response, activated by the alternative sigma factor σ^B^, is controlled by a complex regulatory network that integrates diverse stress and starvation signals, including heat [11]. In addition, Spx is a central regulator of the heat and thiol stress response, which is important for the development of thermotolerance. Spx activates the expression of many genes of the heat shock response, including *clpX* and the oxidative stress response e.g. thioredoxin [5,12,13]. Interestingly, Spx can also mediate the inhibition of cell growth by the concurrent transcriptional down-regulation of many translation-related genes [14].

Another fast acting bacterial stress response system is the stringent response (SR), which is mediated by the second messenger alarmones (p)ppGpp [15]. The synthesis and hydrolysis of (p)ppGpp is catalyzed by RelA/SpoT homologs (RSH) which contain within the N-terminal part synthetase and hydrolase domains (bifunctional Rel or SpoT subgroup) or an active synthetase and an inactive hydrolase domain (RelA subgroup) together with additional regulatory domains at the C-terminus [16]. RSH can therefore direct both synthesis and, in the case of Rel, hydrolysis of (p)ppGpp. The enzyme activity of RelA or Rel is stimulated by association with uncharged tRNAs with the ribosome, thereby mediating (p)ppGpp synthesis upon amino acid starvation [17–21]. In addition to this long multidomain RSH form, monofunctional small alarmone synthetases (SAS) or small alarmone hydrolases (SAH) with single synthetase or hydrolase domains are present in many bacteria [16]. In *B. subtilis*, alarmone levels are controlled by Rel (often referred to as RelA), a bifunctional, RSH-type synthetase/hydrolase as well as two SAS proteins [22, 23].

The synthesis and hydrolysis of (p)ppGpp allows the activation or repression of different cellular pathways by modulating various enzyme activities involved in GTP homeostasis, replication, transcription and translation, not only in response to amino acid starvation, but also to various other signals or stresses. It was observed for different bacteria that additional and diverse starvation or stress signals can activate the SR via interacting proteins or metabolites that bind and modulate the activity of RSH-type enzymes, or by transcriptional or post-translational regulation of monofunctional SAS [24, 25]. *B. subtilis* and related Firmicutes lack a DksA homolog and a direct binding site for (p)ppGpp on RNA polymerase (RNAP) which mediate positive and negative stringent regulation in *E. coli* and other proteobacteria. Instead, in *B. subtilis* (p)ppGpp can exert transcriptional regulation via a drop in GTP levels caused by the direct inhibition of multiple enzymes of the GTP synthesis pathway [26, 27]. Thereby, transcription of ribosomal RNA (rRNA) and ribosomal protein (r-protein) genes from promoters that initiate transcription with GTP is strongly reduced, while in turn promoters that initiate with ATP are activated [28, 29]. In addition, the global regulator CodY is regulated by GTP via an allosteric binding site and de-represses amino acid biosynthesis genes and other pathways during the SR [30]. Beyond regulation of transcription, (p)ppGpp can inhibit translation initiation and elongation by binding, for example, to the translation initiation factor IF-2 and other ribosome-associated GTPases [31–33]. With its ability to inhibit translation and growth, the SR was also implied in persister cell formation and development of antibiotic tolerance [34]. In addition, virulence as well as survival of pathogens during infection was strongly affected in *rel* and (p)ppGpp^0^ mutant strains [15, 35].

During exposure to heat and oxidative stress, we and others previously observed in *B. subtilis* a pronounced down-regulation of rRNA and r-protein genes that resembled the pattern of the SR [13,14,39,40]. Thus, we hypothesized that the alarmone (p)ppGpp and the SR-like response could be part of the heat shock response of *B. subtilis*. Therefore, we investigated the role of the SR and its intricate and mutual involvement with the cellular stress response during various proteotoxic stress conditions, including heat shock conditions, such as growth at high temperatures (50 °C), or thermoresistance and thermotolerance development [5, 41].

Consistent with our hypothesis, we show here that the cellular level of (p)ppGpp was increased upon heat shock and also upon salt and oxidative stress. In addition, artificially raised alarmone levels conferred increased stress tolerance and a (p)ppGpp^0^ strain appeared more stress sensitive. The presence of the bifunctional Rel was necessary and sufficient for the observed stress induced increase of (p)ppGpp. We analyzed changes in the transcriptome with RNA-sequencing (RNA-seq) experiments of wildtype, *rel* and (p)ppGpp^0^ *B. subtilis* strains at raised temperatures and observed pleiotropic adjustments of transcription typical for SR, which was heat-dependent but also partially influenced by (p)ppGpp levels. However, the presence or absence of this second messenger had a more significant and immediate impact on limiting the translation of heat shocked cells. Our results suggest a model in which (p)ppGpp and Spx appear to play a complementary and partially redundant role in stress-mediated readjusting of transcription. In addition, we observed a prominent and instantaneous effect of the cellular alarmone (p)ppGpp levels on limiting translation, allowing the fast reallocation of cellular resources by raising the cellular repair capacity and concurrently reducing the protein load on the PQS during stress.

## Results

### Regimes for monitoring of heat shock stress response in *B. subtilis*

In this study, we monitored the stress response of *B. subtilis* by application of different, but related, heat shock conditions: (i) growth and heat shock at 50°C, a temperature that is non-lethal in *B. subtilis* but already induces a significant heat shock response with a raised expression of chaperones and proteases, (ii) resistance to severe heat shock by measuring the survival of exponentially growing cells exposed to a severe, lethal heat shock at 53 °C, which can also be considered as thermoresistance (37/53°C) (Fig. 1A), and (iii) the development of thermotolerance by measuring the survival of exponentially growing cells primed by a 15 min mild pre-shock at 48 °C before their exposure to the severe heat shock at 53 °C (48/53°C) (Fig. 1A). We experimentally established that 55°C was an appropriate temperature to examine the impact of severe heat on *B. subtilis* cells growing on agar plates. In addition to exposure to these various heat conditions, we also examined other potentially proteotoxic stresses, such as salt and oxidative stress [5,14,41].

**Fig. 1:**
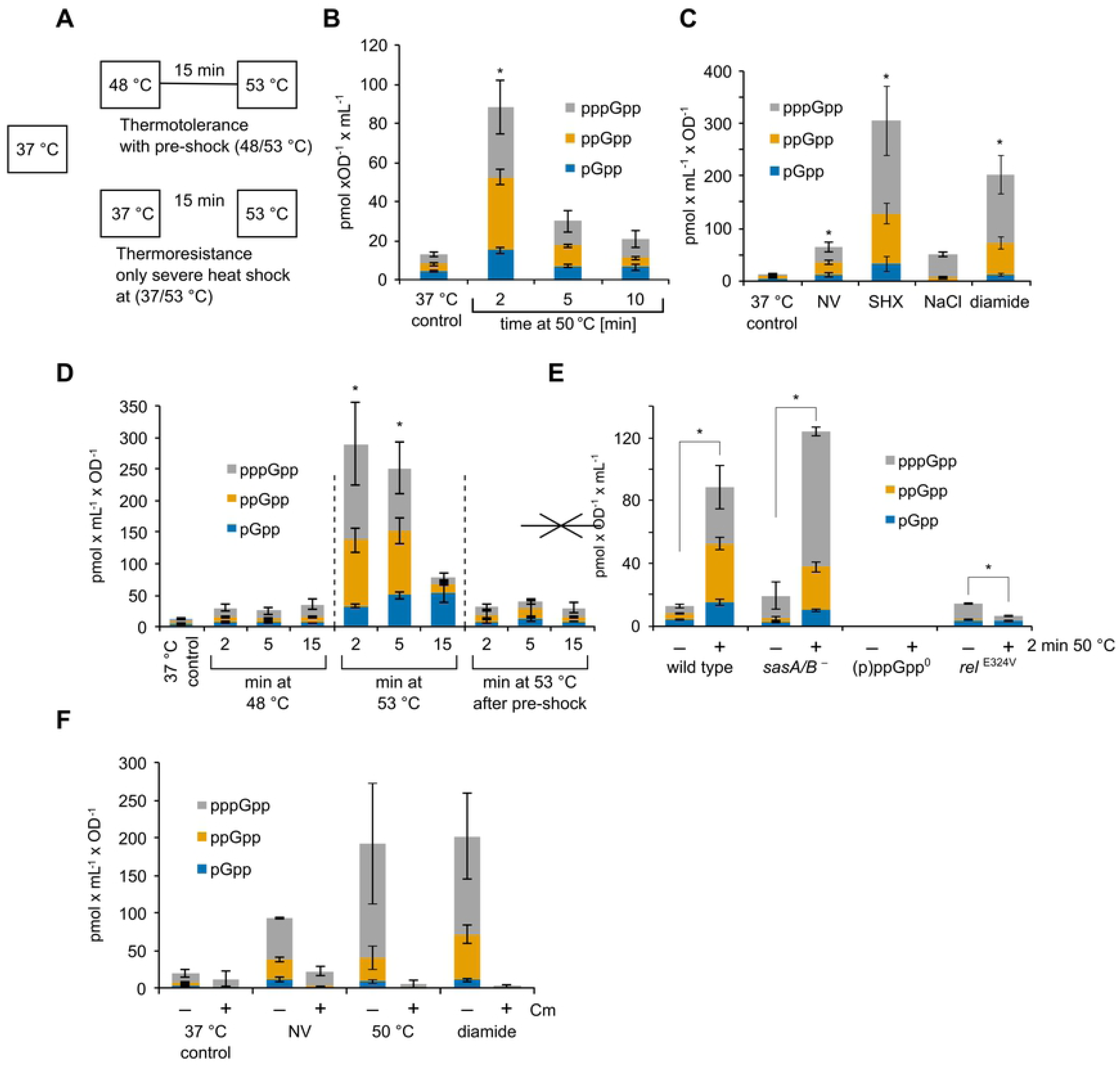
(p)ppGpp levels are increased by heat shock and stress. **(A)** Outline of the thermotolerance protocol. A culture of cells growing exponentially at 37 °C is divided and incubated at 48 °C or left at 37 °C. After 15 min, both cultures are shifted to 53 °C. **(B-F)** Levels of pGpp, ppGpp and pppGpp under different conditions. **(B)** Cells were grown in minimal medium to OD_600_ of 0.4 and transferred to 50 °C or treated with 0.5 mg ml^−1^ DL-norvaline for 10 min. Means and and standard error of mean (SEM) of four independent experiments are shown. Asterisks (*) indicate significance (*p_adj._* ≤ 0.05) of combined pGpp, ppGpp and pppGpp levels according to the Kruskal-Wallis and Dunn-Bonferrroni test**. (C)** Cells were grown in minimal medium to the mid-exponential phase (OD_600_ ∼ 0.4) and treated with DL-norvaline (NV; 0.5 mg ml^−1^), serine hydroxamate (SHX; 5 µg ml^−1^), NaCl (6 %) or diamide (0.5 mM) for 10 min. Means and SEM of three to four independent experiments are shown. Asterisks (*) indicate significance (*p_adj._* ≤ 0.05) of combined pGpp, ppGpp and pppGpp levels according to the Kruskal-Wallis and Dunn-Bonferrroni test. **(D)** Wildtype cells were grown at 37 °C and shifted to 48 °C for 15 min (pre-shock), then to 53 °C or directly to 53 °C. Samples were taken at 2, 5 and 15 min. Means and SEM of four independent experiments are shown. Asterisks (*) indicate significance (*p_adj._* ≤ 0.05) of combined pGpp, ppGpp and pppGpp levels according to the Kruskal-Wallis and Dunn-Bonferrroni test. **(E)** Wildtype cells or strains with mutations in (p)ppGpp synthetases (*sasA/B*^−^: BHS204, *rel*^E324V^: BHS709; (p)ppGpp^0^: BHS214) were treated with or without heat shock at 50 °C for 2 min. Means and SEM of three to six independent experiments are shown. No alarmone peaks were detected in the (p)ppGpp^0^ mutant (lower limit of quantification: 0.26 pmol × mL^−1^ × OD^−1^).. Asterisks (*) indicate significant changes (*p* ≤ 0.05) of combined pGpp, ppGpp and pppGpp levels according to Welch’s *t-*test. **(F)** The influence of chloramphenicol on alarmone accumulation during stress. Cells were grown in minimal medium and treated with DL-norvaline (0.5 mg ml^−1^) for 10 min, heat shock at 50 °C for 2 min or diamide (1 mM) for 10 min. Chloramphenicol (Cm, 25 µg ml^−1^) was added at the same time to one part. Means and SEM of two independent experiments are shown.

### Cellular (p)ppGpp levels increase during heat shock exposure

To investigate the impact of heat on the stringent response, we first assessed the intracellular levels of the alarmones pGpp, ppGpp and pppGpp during the heat shock response at 50 °C. To do so, cells were grown at 37 °C in minimal medium to an optical density at 600 nm (OD_600nm_) of 0.4, and subsequently they were treated with a single, non-lethal temperature upshift to 50 °C in order to induce the heat shock response. After 2, 5 and 10 minutes of incubation at 50 °C, the intracellular levels of the three alarmones (i.e. pGpp, ppGpp and pppGpp) were examined by liquid chromatography (LC)-coupled mass spectrometry (LC-MS). Already after 2 minutes, the alarmone levels increased approx. seven-fold (from 13 to 88 pmol OD^−1^ ml^−1^) (Fig. 1B). The observed alarmone accumulation after 2 minutes at 50 °C was in a similar range to that previously observed upon amino acid starvation induced by DL-norvaline, serine hydroxamate, salt stress induced by 6 % (w/v) NaCl or 0.5 mM diamide, a strong oxidant of thiol groups (Fig. 1 C) [39, 42]. It should be noted that the (p)ppGpp levels increased only transiently during heat shock and reduced to almost basal levels after approximately 10 minutes (Fig. 1B). Thus, we conclude that exposure to a non-lethal heat shock at 50°C elicits a fast, but transient, increase of the alarmones pGpp, ppGpp and pppGpp.

Having shown that (p)ppGpp levels transiently increase during heat shock, we next assessed the levels of the alarmones under thermoresistance conditions (37/53°C), after priming (37/48°C), as well as under thermotolerance conditions (48/53°C) (Fig. 1A & D). When we examined (p)ppGpp levels upon those temperature shifts, we observed transiently increased (p)ppGpp levels (Fig. 1D). The alarmone levels were particularly high during the severe heat shock shift at 37/53 °C (about 25-fold increase) and the induction was lower both for a 37/ 48°C or 48/53°C shift (about 2-3 fold increase) (Fig. 1D). Thermotolerant cells that were exposed to 48/53 °C showed a comparable alarmone level to cells exposed to 48 °C or 50 °C after 5 min (2-3 fold), while cells only exposed to a higher lethal heat shock of 37/53 °C display a relative much higher alarmone level (Fig. 1B,C). The primed thermotolerant cells appear to be able to somehow limit the alarmone synthesis, when exposed to the lethal heat shock.

The synthesis of (p)ppGpp that occurs during activation of the SR is normally accompanied by a fast reduction of cellular GTP levels in cells treated with serine hydroxamate (SHX) or DL-norvaline (NV) [27] and also after exposure to salt or diamide (Fig. S1A). Therefore, we were also interested in monitoring changes in GTP levels under conditions of heat shock but interestingly we do not observe a reduction in GTP levels after exposure to 50 °C (Fig. S1A). Notably, the GTP levels were at a comparable high level (FigS1B) during temperature upshifts of 37/48 °C, 37/53 °C and 48/53 °C, however GTP levels appeared a little lower for all temperature upshifts after 15 min incubation (Fig. S1B).

Taken together, we show that exposure to heat shock elicits a fast, but transient, increase of the alarmones pGpp, ppGpp and pppGpp, while not immediately affecting the GTP levels. Therefore, it seems that alarmone levels exhibit a graded response to stress, which appears to correlate to the temperature levels and possibly the heat stress intensity the cells are exposed to.

### Rel is the main source for (p)ppGpp synthesis during stress response

Next, we aimed to identify the major source of (p)ppGpp during the heat stress response. To this end, strains with mutations that disrupt the (p)ppGpp synthetase activity of the proteins encoded by either *sasA/ywaC* and *sasB/yjbM* (*sasA/B*^−^ strain) or *rel* (*rel*^E324V^; inactive synthetase) were assayed for (p)ppGpp accumulation and GTP levels upon heat shock at 50 °C for 2 min (Fig. 1E, S1C). As a control, (p)ppGpp accumulation was also measured in a (p)ppGpp^0^ strain bearing inactivating mutations in all three alarmone synthetase genes (*sasA*, *sasB* and *rel*) (Fig 1E). In addition to monitoring (p)ppGpp accumulation directly, the (p)ppGpp-dependent transcription of *hpf* was employed as an additional read-out for the activation of the stringent response (Fig. S1D) [43, 44]. As expected, alarmone nucleotides were not detected in the (p)ppGpp^0^ mutant under any conditions, neither stress or non-stress, consistent with finding that Rel, SasA and SasB are the only sources of (p)ppGpp in *B. subtilis* [23] (Fig. 1E). We observed that the *sasA*/*B^−^* strain also exhibited accumulation of (p)ppGpp (Fig. 1E) and up-regulation of the *hpf* transcript similar to the wildtype *B. subtilis* cells upon heat exposure (Fig. S1D), indicating that the activity of SasA and SasB is dispensable for (p)ppGpp production during heat stress. By contrast, the *rel*^E324V^ strain accumulated negligible amounts of (p)ppGpp in response to heat, with the levels even dropping after heat shock (Fig. 1E). Consistently, up-regulation of the *hpf* transcript and accumulation of the Hpf protein in response to stress was also strongly impaired in the *rel*^E324V^ strain (Fig. S1D, S12D). Together, these results strongly suggest that Rel is the main source of (p)ppGpp during heat stress.

Activation of Rel during amino acid starvation requires the presence of uncharged tRNA on the ribosome [17, 18]. A first indication of such a connection between SR and Rel activation in conjunction with the ribosome was the initial observation that (p)ppGpp accumulation upon starvation for amino acids was almost completely suppressed in the presence of the translation-inhibitor chloramphenicol [45]. To probe, whether Rel activation during heat or oxidative stress could utilize a similar pathway, we measured alarmone levels in stressed cells in the presence or absence of chloramphenicol (Fig. 1F, S1E). Interestingly, the addition of chloramphenicol completely suppressed alarmone accumulation and resulted in increased GTP levels upon heat and diamide treatment. Notably, chloramphenicol treatment of unstressed cells did not induce a SR, but decreased the basal (p)ppGpp levels and slightly increased GTP (Fig. 1F, S1E). These observations indicate that heat and oxidative stress could activate Rel in a similar manner to each other and similar to the pathway suggested for amino acid starvation.

### *B. subtilis* cells lacking the alarmone are more sensitive to stress

To assess the importance of alarmone production for cellular survival under heat stress, we monitored growth of the wildtype, (p)ppGpp^0^, *sasA/B^−^* and *rel*^E324V^ strains at 37 °C and 55 °C (Fig. 2A, B). As expected, no obvious growth defects were observed for any of the strains at 37 °C. While the cellular survival of the *sasA/B^−^* strain at 55 °C was identical to that of the wildtype strain, strong growth defects were evident for the (p)ppGpp^0^ and *rel*^E324V^ strains at 55 °C. These findings suggested that production of (p)ppGpp by Rel, but not SasA/B, is critical for survival of *B. subtilis* cells under heat stress. This prompted us to also investigate whether production of (p)ppGpp by Rel is critical for survival of *B. subtilis* cells under other stress conditions, such as high salt or oxidative stress. Indeed, severe growth defects were observed for both the (p)ppGpp^0^ and *rel*^E324V^ strains under oxidative heat and salt stress, whereas the growth behavior of the *sasA/B^−^* strain again resembled the wildtype strain under the same conditions (Fig. 2C, D). Collectively, these findings suggest that production of (p)ppGpp by Rel is critical for survival of *B. subtilis* cells, not only under heat stress, but also conditions of oxidative and salt stress.

**Fig. 2:**
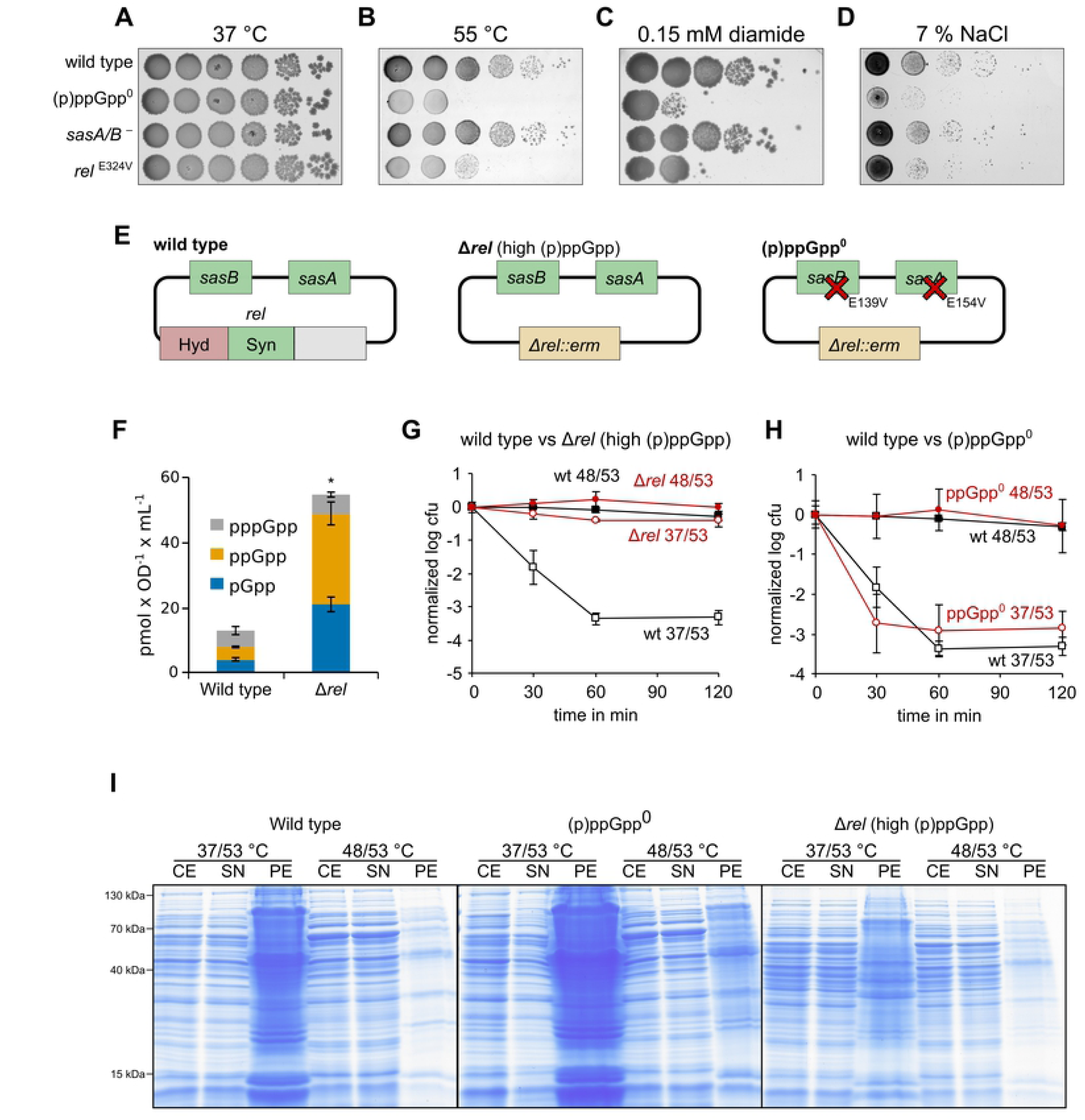
Increased (p)ppGpp levels confer high heat stress resistance. **(A-D)** Growth of strains with mutations in (p)ppGpp synthetases (*sasA/B*^−^: BHS204, *rel*^E324V^: BHS709; (p)ppGpp^0^: BHS214) on agar plates at 37 °C, during heat stress (55 °C), oxidative stress (0.1 mM diamide) or salt stress (total concentration of 7 % (w/v) NaCl) over night. **(E)** Outline of the genotypes and the (p)ppGpp synthesis capabilities of the assessed wildtype, Δ*rel* (BHS126 and BHS368) and (p)ppGpp^0^ (BHS214 and BHS319) strains. **(F)** Cellular alarmone levels of wildtype and Δ*rel* strains. Asterisks indicate significant changes (*p* ≤ 0.05) of combined pGpp, ppGpp and pppGpp levels according to Welch’s *t-*test. Means and SEM of three independent experiments are shown. **(G/H)** Thermotolerance and survival of wildtype (black lines) and mutant strains (red lines) at 53 °C. Means and SEM of at least three independent experiments are shown. Open symbols: no pre-shock, closed symbols: 15 min pre-shock at 48 °C. **(I)** Accumulation of protein aggregates during heat stress at 53 °C without (37/53 °C) or with (48/53 °C) pre-shock. CE: cell extract, SN: supernatant, PE: pellet (aggregated protein fraction).

### High cellular (p)ppGpp levels confer elevated heat stress resistance

Next, we asked whether (p)ppGpp alarmone levels influence thermotolerance development and survival. To do this, we utilized the (p)ppGpp^0^ strain, which cannot synthesize (p)ppGpp (Fig. 1E) as well as a *rel* deletion strain that displays raised (p)ppGpp (Fig. 2F) and lowered GTP levels (Fig. S2A). The high (p)ppGpp levels in the *rel* deletion strain arise because Rel is the only alarmone hydrolase in *B. subtilis* and causes an overall decrease in growth rate (Fig. 2F, S2A, B), as reported previously [23, 46]. For completeness, we also assayed *sasA* and *sasB* deletion strains. As expected, exposure of wildtype *B. subtilis* cells to heat shock at 37/53°C led to a dramatic reduction in survival, e.g. 1000-fold (3-log) reduction in viability with 60 min heat shock at 53 °C, whereas survival remained unaltered when cells received a pre-shock at 48 °C for 15 min before being exposed to the lethal heat shock at 53 °C (Fig. 2G). Similarly, *B. subtilis* strains with single deletions in *sasA* or *sasB* phenocopied the wildtype strain for thermotolerance development (Fig. S2C-D), as they did for heat shock resistance (Fig. 2A-B and Fig. S2E). By contrast, we observed that *rel* deletion resulted in strongly increased thermoresistance, which was apparent from the high number of cells still able to form colonies during the otherwise lethal heat shock (Fig 2G). Consistently, we also observed a strong reduction in protein aggregation during the 37/53 °C heat shock (Fig 2 I). While no significant effect on thermotolerance development was observed in the (p)ppGpp^0^ strain (Fig. 2 H), the (p)ppGpp^0^ strain exhibited more protein aggregation when exposed to 37/53 °C heat shock (Fig 2 I).

To confirm that the elevated heat resistance phenotype of the *rel* strain was caused by the elevated levels of the alarmone (p)ppGpp, rather than the absence of the Rel protein, we expressed a truncated form of the *E. coli* RelA (*RelA_hyper_*) that exhibits constitutive and hyperactive alarmone synthetase activity *in trans* in wildtype *B. subtilis* cells [47, 48]. As a control, we also expressed a truncated form of the *E. coli* RelA (*RelA_inactive_*) that has no alarmone synthetase activity [47, 48]. In a second approach, we examined the (p)ppGpp^0^ strain expressing *in trans B. subtilis* Rel with mutations that inactivate either the synthetase (Rel^E324V^) or hydrolase (Rel^H77AD78A^) domains. Expression of RelA_hyper_ or hydrolase-inactive Rel^H77AD78A^ resulted in increased alarmone levels (Fig. S3A) and conferred high thermoresistance (Fig. S3B,C), as observed for the Δ*rel* strain (Fig. 2G). By contrast, strains expressing RelA_inactive_ or the synthetase-inactive Rel^E324V^ did not display increased alarmone levels (Fig. 3E), nor increased survival to severe heat stress (Fig. S3D,E).

**Fig. 3:**
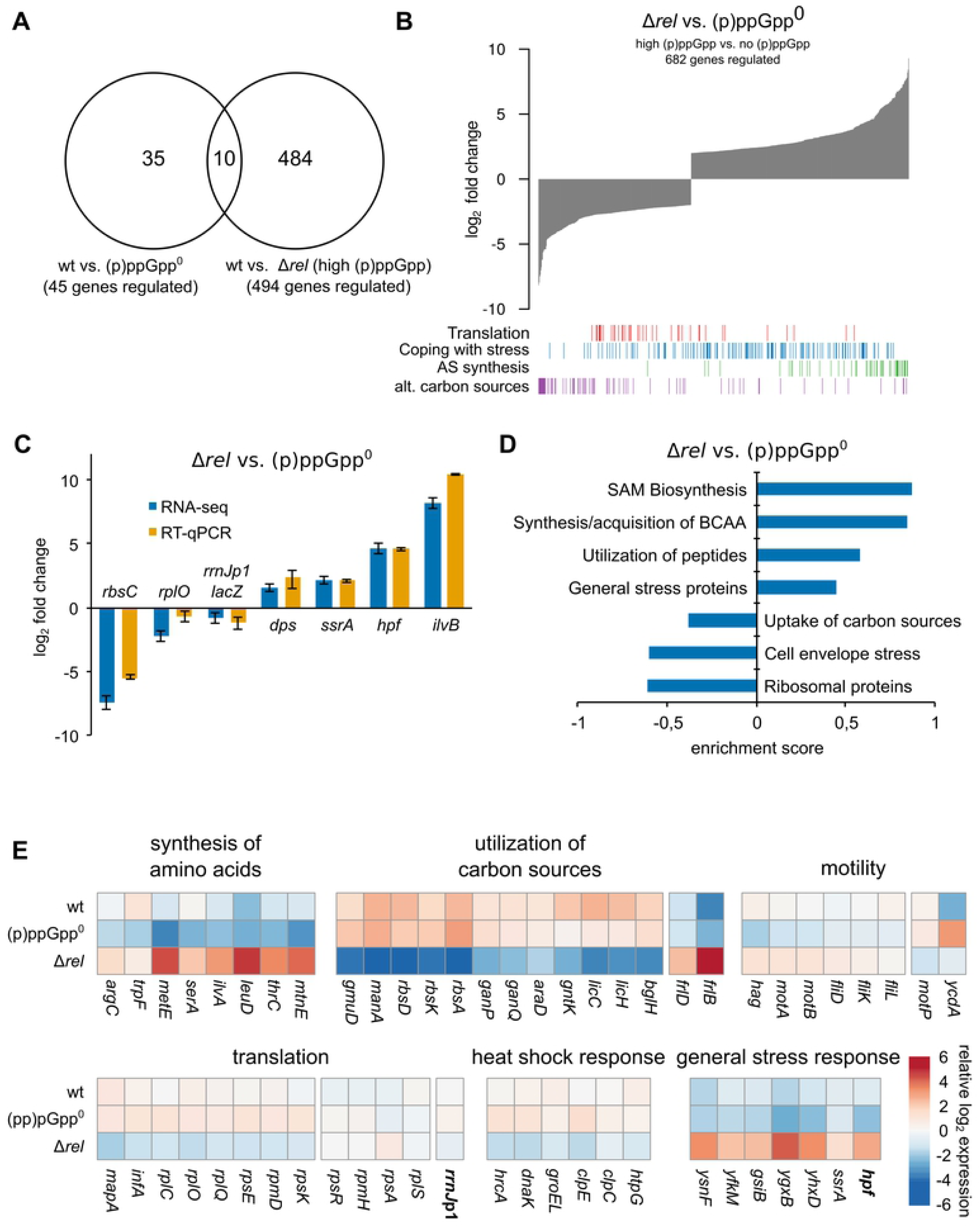
(p)ppGpp-mediated global changes in the transcriptome. **(A)** Venn diagram showing the number of significantly regulated genes in Δ*rel* or (p)ppGpp^0^ strains vs. wildtype. **(B)** Global differences in gene expression in Δ*rel* versus (p)ppGpp^0^ strains. Bar tracks indicate the distribution of genes in the respective functional groups. **(C)** Comparison of the relative transcription changes of selected genes in, Δ*rel* and (p)ppGpp^0^ strains during exponential growth at 37 °C as determined by RNA-seq or RT-qPCR from independent experiments. Means and SEM of three replicates are shown. **(D)** Selected category results of the gene set enrichment analysis from regulated transcripts in Δ*rel* vs. (p)ppGpp^0^ cells. Positive/negative enrichment scores represent enrichment in the up- or down-regulated genes. **(E)** Heatmap showing the expression changes of selected transcripts in wildtype, (p)ppGpp^0^ or Δ*rel* strains. Values represent normalized log_2_ scaled read counts centered on the mean expression level of each transcript.

High (p)ppGpp levels during the SR lead to a decrease in cellular GTP levels and this decrease is known to be intricately involved in causing the transcriptional changes during SR [27, 28] (Fig. S1, S2A, S3A). To examine, whether the resistance to heat stress observed in the Δ*rel* strain could be mediated simply by lowering cellular GTP levels, wildtype cells were treated with decoyinine, an inhibitor of GMP synthetase, which results in a significant drop of cellular GTP levels (> 3-fold) without increasing (p)ppGpp levels [49, 50]. Treatment with 250 or 400 µg ml^−1^ decoyinine resulted only in moderately increased thermoresistance and moderately decreased thermotolerance (Fig. S4). However, we could not observe the strongly increased thermoresistance as we observed before in the presence of raised (p)ppGpp levels (Fig 2 F, G, Fig S3). In addition, higher decoyinine concentrations (1000 µg ml^−1^) even abolished both thermoresistance and thermotolerance development (Fig S4). These experiments suggest that lowered cellular GTP levels, which turn the transcriptional stringent response on [27,28,51], is not sufficient to elicit heat resistance as observed in strains with elevated (p)ppGpp levels (Fig 2 F, G, Fig S3).

From these observations we infer that raised (p)ppGpp levels are sufficient to confer increased stress resistance and reduced levels of heat-induced protein aggregates. This phenotype is dependent on the levels of alarmones and not the presence or absence of the specific Rel protein *per se*, since it could be reconstituted by *in trans* expression of full-length or truncated Rel or RelA protein variants from *B. subtilis* or *E. coli* that actively synthesized (p)ppGpp (Fig. 2, S3). The SR mediated drop in cellular GTP levels was not observed during heat shock response (Fig. S1) and an artificial reduction of cellular GTP levels had only a moderate effect on thermoresistance and even abolished thermotolerance (Fig. S4). Taken together, these experiments suggest that the cellular level of the second messenger (p)ppGpp *per se* appears to be important for the modulation and enhancement of the heat shock response in *B. subtilis* cells, since a strain lacking the alarmone is more stress sensitive (Fig. 2 A-D, H, I) and strains with constitutively raised alarmone levels are much more stress resistant (Fig. 2 F, G, I).

### Constitutive stringent response in Δ*rel* cells results in global transcriptional changes

To obtain further insights into the impact of *rel* deletion and (p)ppGpp accumulation on transcriptome changes, we performed RNA-seq analyses and annotated transcription start sites (TSS) of exponentially growing wildtype, (p)ppGpp^0^ and Δ*rel* strains (Fig. S5, see S1 Text for a detailed analysis, Dataset S1, Dataset S2, Dataset S3). Since down-regulation of “stable” rRNA is a hallmark of the SR, we introduced a previously established chromosomal *rrnJ*p1-*lacZ* fusion into the assessed strains, thereby allowing us to follow the activity of this rRNA promoter using the *lacZ* reporter [14]. Only small changes between wildtype and (p)ppGpp^0^ strains (45 genes significantly regulated, Fig. 3A, Fig. S6B) were observed during non-stressed growth, while Δ*rel* cells exhibit broad transcriptional changes compared to wildtype cells (494 genes regulated, Fig. 3A & Fig. S6C). However, the full extent of the impact of (p)ppGpp was revealed when we compared the transcriptome of the (p)ppGpp^0^ with Δ*rel* strain (Fig. 3A, B). Here we observed a differential regulation of the expression of 682 genes with a broad down-regulation of translation-related genes known to be part of the SR regulon (Fig. 3B). We performed RT-qPCR experiments with independent samples of the (p)ppGpp^0^ and Δ*rel* strains, measuring the transcripts of selected genes known to be under stringent control and observed a good correlation with the RNA-seq data (Fig. 3C). The *rrnJ*p1*-lacZ* transcript was down-regulated 1.7-fold in the RNA-seq experiment and confirmed by RT-qPCR (Fig. 3C). Furthermore, we noticed an extensive up-regulation of genes that function in amino acid synthesis, indicating a de-repression of the CodY regulon (e.g. *ilvB* 294-fold up-regulated, Fig. 3B,D,E, S8, Dataset S2) [27, 52]. Interestingly, we detected a strong decrease in the transcription of CcpA-regulated genes required for the utilization of alternative carbon sources (e.g. *rbsC* 181-fold down-regulated) and a broad regulation of stress-related genes accompanied by an activation of the SigB regulon (e.g. *dps* 2.8-fold up-regulated, *ssrA* 4.5 fold up-regulated). We also observed a reduced transcription of genes regulated by the regulators HrcA and CtsR (e.g. *dnaK* 6.4-fold down-regulated*, clpE* 7.1-fold down-regulated) (Fig. 3C, D, E and Fig. S7, S8, Dataset S2), when comparing the (p)ppGpp^0^ and Δ*rel B. subtilis* strains at 37 °C without heat exposure. Notably, the transcription of *hpf* (*yvyD*), encoding the hibernation promoting factor Hpf, was induced by raised (p)ppGpp levels (24-fold up-regulated, Fig. 3C,E, Fig. S8), confirming that the increased transcription of *hpf* can be considered as a reporter for the activation of the SR [43, 44].

### (p)ppGpp modulates transcription during heat stress response

To study the impact of the SR on the transcriptome during heat exposure, we examined the (p)ppGpp^0^ and Δ*rel* strains not only at 37 °C (Fig. 3), but also at 48 °C, in the same RNA-seq experiment that was used to investigate the thermoresistance (37/53 °C) and thermotolerance (48°/53 °C) conditions (Fig.4 & Fig. 1A) [5, 14]. Thermotolerant cells (48/53 °C) exhibited a pronounced up-regulation of the heat-specific stress response (median 3.4-fold up-regulated) or general stress response (median 3.0-fold up-regulated) as well as comprehensive down-regulation of translation-related genes (median 2.6-fold down-regulated, Fig. 4A, B, S7, Dataset S2) that was, to a lesser extent, also observed in the mild pre-shock (48 °C, median 1.3-fold down-regulated) and severe heat shock (37/53 °C, median 1.2-fold down-regulated) conditions (Fig. S6 A, S7) in agreement with previous observations [14]. The transcription of translation-related genes was generally lower than wildtype levels in Δ*rel* cells (median 1.6-fold down-regulated) and higher in (p)ppGpp^0^ strains under non-stress conditions (37 °C) (Fig. 4C, S7). We could confirm by RT-qPCR that down-regulation of *rrnJ*p1-*lacZ* is partially (p)ppGpp-dependent during thermotolerance development (Fig. 4C,D), which became even more pronounced when the 50 °C heat shock condition was examined (Fig. S8) [14]. Nevertheless, the transcription of the genes encoding conserved chaperones and proteases of the heat shock response were strongly up-regulated upon all temperature up-shifts, independently of the presence or absence of (p)ppGpp (Fig. 4 C). Interestingly, additional qPCR experiments applying a 50 °C heat shock revealed that the heat-induced expression of some SigB regulated genes was impaired in the (p)ppGpp^0^ background, e.g. of *ssrA* (approx. 2-fold lower expression in (p)ppGpp^0^ cells at 50 °C) and *dps* (approx. 3-fold lower expression), indicating a functional connection between the SR and the general stress response (Fig. S8) [53, 54]. However, the majority of genes of the SigB regulon were found to be induced in the (p)ppGpp^0^ strain similarly to wildtype cells at 48 °C (Fig. S7). Likewise, while CcpA-regulated genes were repressed in wildtype and (p)ppGpp^0^ cells under heat shock conditions (Fig. 3B,D, Fig. 4A-C, S7), some genes (e.g. *rbsD, ganP, licH*) were less down-regulated or even induced at 48 °C in the (p)ppGpp^0^ strain (Fig. 4C). In contrast, motility-genes were particularly strongly down-regulated by heat in the (p)ppGpp^0^ mutant (median 3.0-fold change, Fig. 4C, S6B, S7), while the down-regulation of these genes appeared to be not significant in wildtype cells at 48 °C (median 1.14-fold change, S7) [55]. Notably, the expression of the *hpf* and *ilvB* transcripts, the transcription of which is positively regulated by the SR, was lower during heat stress in the (p)ppGpp^0^ strain compared to wildtype cells (*hpf*: 3.7-fold lower in (p)ppGpp^0^ cells relative to wildtype at 50 °C, *ilvB*: 1.6-fold lower, Fig. 4D, S8).

**Fig. 4:**
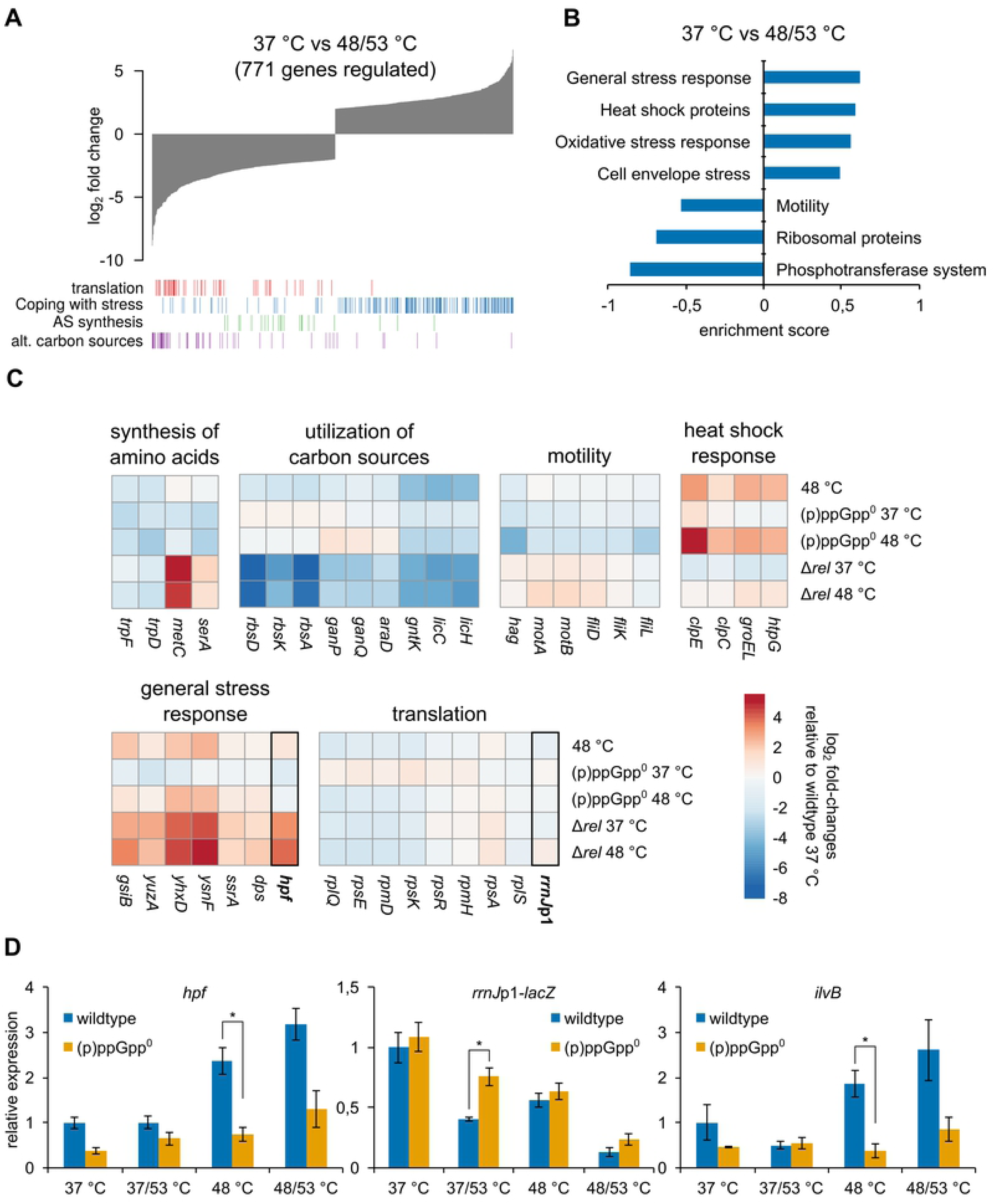
(p)ppGpp mediated transcriptional changes during heat stress. **(A)** Global differences in gene expression of heat shocked, thermotolerant cells (48/53 °C) versus untreated cells (37 °C). Bar tracks indicate the distribution of genes in the respective functional groups. **(B)** Selected category results of the gene set enrichment analysis from regulated transcripts in heat shocked (48/53 °C) cells. Positive/negative enrichment scores represent enrichment in the up- or down-regulated genes. **(C)** Heatmap showing expression changes of selected transcripts during mild heat stress in wildtype, (p)ppGpp^0^ or Δ*rel* cells. Values represent log_2_ fold changes of transcript levels relative to wildtype cells at 37 °C. **(D)** Relative changes in the transcription of selected genes during heat shock in wildtype and (p)ppGpp^0^ strains determined by RT-qPCR. Means and SEM of three replicates are shown. Asterisks indicate significance (*p* ≤ 0.05) according to Welch’s *t-*test.

### Spx and the stringent response act complementary during heat shock

Previously, we reported that Spx, a central regulator of the heat and oxidative stress response, can down-regulate the transcription of translation-related genes and rRNA (ref). However, an *spx* deletion was not impaired in the heat-mediated down-regulation of these genes [14]. Here, we noticed a complex, but clearly detectable, involvement of the SR in the down-regulation of specific genes during heat stress (Fig. 4C-D), suggesting an intricate regulation of these genes by different factors, including Spx and (p)ppGpp. To test for such a concurrent and complementary transcriptional regulation, a *B. subtilis* strain combining a *spx* deletion with the (p)ppGpp^0^ mutations was constructed. Strikingly, down-regulation of *rrnJ*p1-*lacZ* upon heat shock was completely abolished in this (p)ppGpp^0^ Δ*spx* strain, indicating a concurrent and complementary activity of both regulators on this promoter (Fig. 5A). However, the transcription of some r-protein genes was also down-regulated in the (p)ppGpp^0^ Δ*spx* strain (Fig. S9A), suggesting additional factors beyond Spx and (p)ppGpp, that can also influence the promoter and/or the stability of these transcripts. Interestingly, this (p)ppGpp^0^ Δ*spx* strain, lacking both regulators, also displayed a slow growth phenotype at 37 °C and a more severe growth defect at 50 °C compared to the strains with single deletions of (p)ppGpp^0^ or Δ*spx* (Fig. 5B, Fig. S9B). This experiment suggests a genetic interaction of the SR and the *spx* regulon under heat stress conditions. In addition, the (p)ppGpp^0^ Δ*spx* strain accumulated more heat-induced protein aggregates at 50 °C than cells lacking either (p)ppGpp or *spx* (Fig. S9 C).

**Fig. 5:**
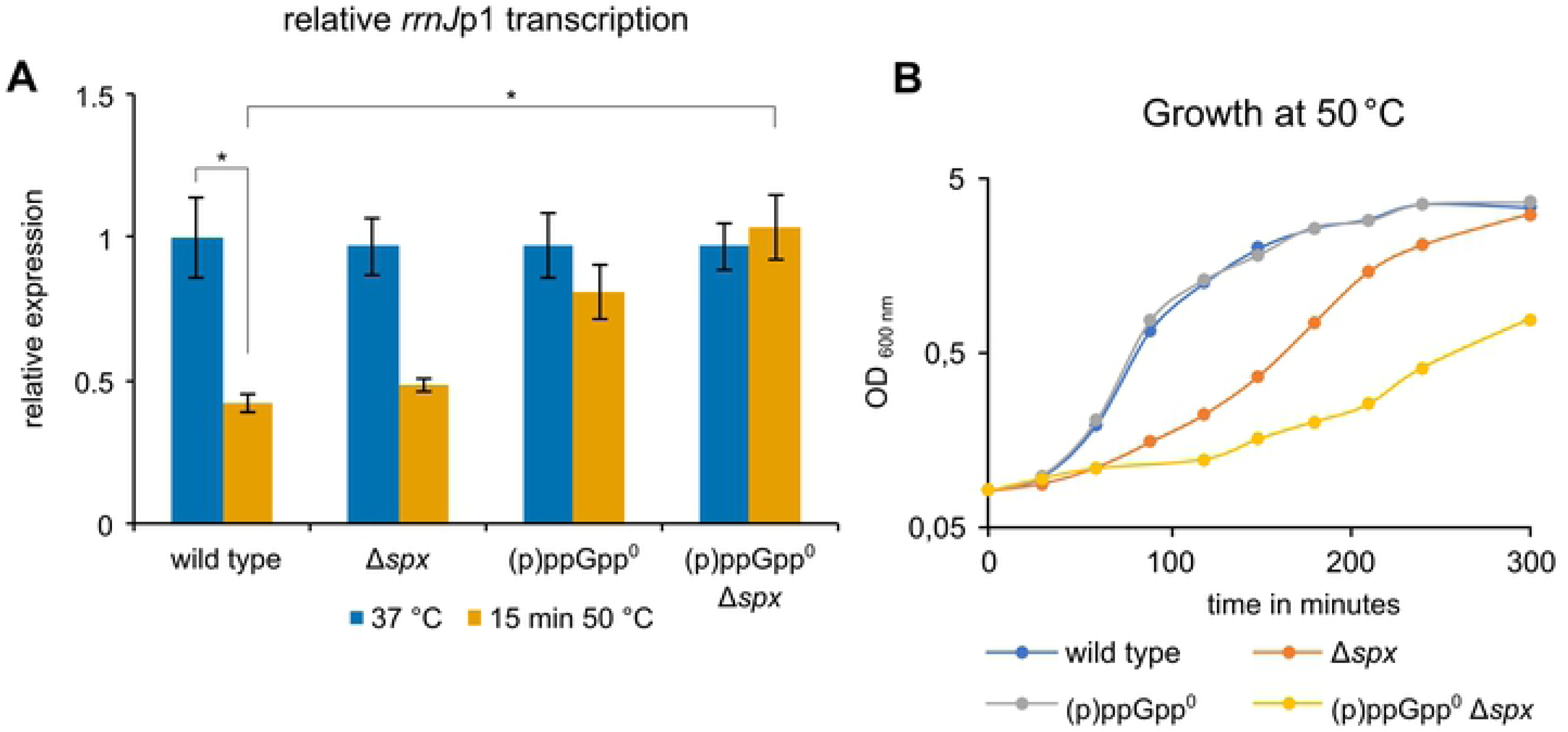
(p)ppGpp and Spx act complementary during heat shock. **(A)** Heat mediated down-regulation *of rrnJ*p1-*lacZ* transcription in wildtype (BHS220), Δ*spx* (BHS222), (p)ppGpp^0^ (BHS319) and Δ*spx* (p)ppGpp^0^ (BHS766) strains as determined by RT-qPCR. Means and SEM of three independent experiments are shown. Asterisks indicate significant changes (*p* ≤ 0.05) of combined pGpp, ppGpp and pppGpp levels according to Welch’s *t-*test. **(B)** Growth of the same strains in LB medium at 50 °C.

When mutations in *rpoA* were introduced in the (p)ppGpp^0^ strain that abolish Spx-mediated up- and down-regulation (*cxs*-1/*rpoA*^Y263C^), or interfere only with Spx-mediated repression of rRNA while still allowing up-regulation of redox chaperones (*cxs*-2 / *rpoA*^V260A^) [14], only the (p)ppGpp^0^ *cxs*-1 strain displayed a severe growth defect as observed for the (p)ppGpp^0^ Δ*spx* strain (Fig. S9B). This experiment suggests that the Spx-mediated up-regulation of stress response genes, and not the ability to down-regulate translation related genes, is required for efficient growth in the (p)ppGpp^0^ background. Notably, (p)ppGpp is sufficient for the down-regulation of translation-related genes during norvaline-induced amino acid limitation, while Spx is not required for this process (Fig. S10A). Conversely, Spx can act on rRNA promoters independently of (p)ppGpp *in vivo* (Fig. S10B) [14]. In addition, *in vitro* transcription experiments with purified Spx and RNAP gave no indications that ppGpp could directly influence Spx transcriptional activation or inhibition of RNAP (Fig. S10C).

Together, these observations suggest that both regulatory systems act concurrently but independently on rRNA and r-protein promoters, allowing the inhibition of transcription of translation-related genes by Spx and to a minor extend also by the (p)ppGpp-mediated transcriptional response.

### (p)ppGpp curbs translation during heat stress

We observed that the raised levels of (p)ppGpp, but not the transcriptional reprogramming during SR, appears to be necessary for the observed strong heat stress resistance (Fig 2, 3, 4, Fig S3 & S4). Therefore, we wanted to determine the impact of (p)ppGpp on translation during heat stress. To this end, a method for pulse-labeling newly synthesized nascent peptide chains using low amounts of puromycin was utilized to estimate protein synthesis rates (see S1 Text, Fig. S11). When we examined growing cells at 50° C, or cells exposed to thermotolerance conditions (48/53 °C), we observed that the Δ*rel* strain always exhibited a lower translation rate compared to the wildtype cells (Fig. 6A,B), consistent with its raised (p)ppGpp levels and the observed “stringent” phenotype of this strain. By contrast, the “relaxed” (p)ppGpp^0^ strain always exhibited higher translation rates (Fig. 6A,B), indicating a more deregulated translation. During the non-lethal 50 °C heat shock, translation rates transiently increased in all strains (Fig. 6A), corresponding with the high growth rate at this temperature (Fig. S9B). Nevertheless, the (p)ppGpp^0^ strain still displayed significantly higher translation rates compared to wildtype and the Δ*rel* strains (Fig. 6A and 6B).

**Fig. 6:**
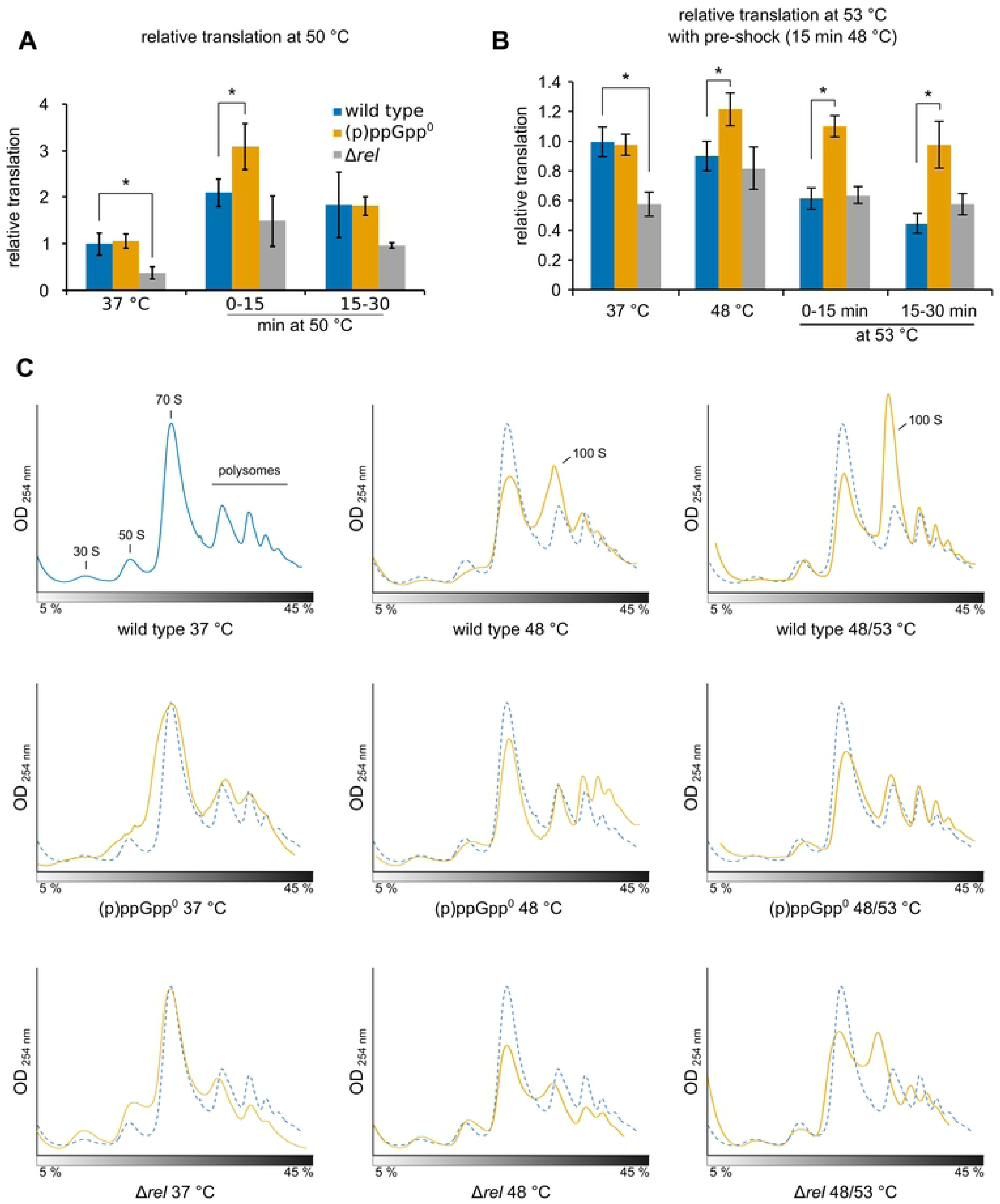
(p)ppGpp modulates translation during stress response. **(A/B)** Relative translation (estimated from puromycin incorporation) of wildtype, (p)ppGpp^0^ (BHS214) and Δ*rel* (BHS126) strains during heat stress **(A)** at 50 °C or **(B)** at 48 °C, 53 °C or 48/53 °C. 1 µg ml^−1^ puromycin was added for 15 min to the medium directly after (0-15 min) or 15 min after shifting the sample to the indicated temperatures. Means and SEM of four independent experiments are shown. Asterisks indicate significance (*p* ≤ 0.05) relative to wildtype according to Welch’s *t-*test. **(C)** Sucrose gradient profiles of extracts from wildtype,(p)ppGpp^0^ (BHS214) or Δ*rel* cells (BHS126) with or without heat shock at 48 °C or 48/53 °C for 15 min each.

Treatment with a lethal temperature shift (37/53 °C) without pre-shock resulted in a strong decrease in translation efficiency in wildtype and (p)ppGpp^0^ strains, whereas translation in Δ*rel* cells was transiently increased (Fig. S12A), in agreement with the observed high heat-resistance of this strain (Fig. 2G). Interestingly, translation was strongly decreased in (p)ppGpp^0^ cells at 37/53 °C, while wildtype cells still maintained active translation under this condition (Fig. S 12A). The lowered translation activity in (p)ppGpp^0^ cells appears to be accompanied by a strong reduction of the levels of cellular 16S rRNA (Fig. S12C), which could indicate a defect in 16S rRNA maturation and the assembly and/or activity of the small ribosomal subunit.

The (p)ppGpp^0^ strain also failed to induce expression of the *hpf* gene during heat stress and did not accumulate the Hpf protein (Fig. 4C,D, S12D). Thus, the formation of 100 S disomes upon heat stress, which was clearly visible in the ribosome profiles of wt and Δ*rel* cells, especially under thermotolerance conditions, was strongly reduced in the (p)ppGpp^0^ strain (Fig. 6C). However, the observed apparent degradation of the 16S rRNA under severe stress conditions was not prevented by *in trans* expression of Hpf (Fig. S12 E) and overexpression of Hpf could not rescue the heat-sensitive phenotype of (p)ppGpp^0^ strains (Fig. S12F). Also, the addition of translation-inhibiting antibiotics could not rescue this phenotype, indicating that inhibition of translation *per se* is not sufficient to protect ribosomes during severe heat stress (Fig. S12G).

The observed influence of (p)ppGpp on translation suggests that the major impact of (p)ppGpp appears not to be its effect on transcription (Fig. 4, 5), but the direct modulation of translation (Fig. 6, S12), possibly by directly interfering with the activity of different translational GTPases [31, 33]. Conversely, Spx appears to act on transcriptional regulation of stress-response and translation-related genes [14]. To assess the relative impact of Spx on translation, we examined the translation rate in a *B. subtilis* strain encoding an inducible gene for the synthesis of a stable Spx^DD^ variant and observed only a 20 % reduction of translation by Spx^DD^ induction (Fig. S9D). This reduction may be an indirect result of the Spx^DD^-mediated repression of rRNA synthesis with the ensuing decreased synthesis of new ribosomes [14].

In summary, these observations indicate that the intracellular (p)ppGpp second messenger can immediately control translation during heat stress and is involved in the protection of ribosomes from damage upon severe heat stress (Fig. 7).

**Fig. 7:**
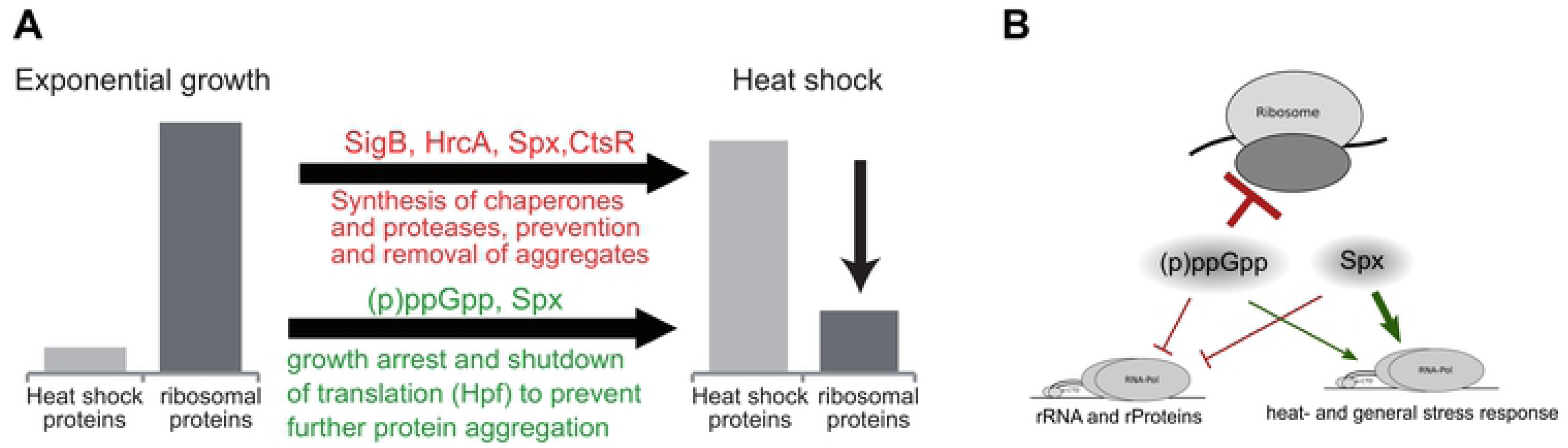
The interplay of the stringent response and the heat shock response. **(A)** Model of the role of the stringent response in the regulatory network of the heat shock response. **(B)** The interplay of Spx activity and the stringent response in the regulation of transcription and translation during the heat shock response.

## Discussion

In this study, we analyzed the role of the SR during heat shock in *B. subtilis*. We could demonstrate that upon heat shock the second messenger (p)ppGpp is rapidly synthesized mostly by Rel and can confer enhanced thermoresistance to these cells. Our data suggests that (p)ppGpp is a pleiotropic regulator, affecting several transcriptional processes, but mostly modulates and protects translation during heat stress. The SR- and Spx-mediated heat shock responses can act concurrently and might be able to complement one another in the down-regulation of rRNA transcription during stress. Overall, our results suggest that limiting translation is an integral part of the *B. subtilis* stress response (Fig. 7).

### The activation of the stringent response during heat stress

The presented results clearly demonstrate a rapid accumulation of (p)ppGpp during heat and other environmental stresses (Fig. 1). In addition, strains unable to synthesize (p)ppGpp are rendered sensitive to high temperatures and accumulate more heat-induced protein aggregates (Fig. 2A-D,H,I). Interestingly, (p)ppGpp synthesis and heat tolerance are solely dependent on the synthetase activity of Rel, indicating that this enzyme is the major contributor of (p)ppGpp under these conditions (Fig. 1E, 2A-D). Activation of the SR by Rel-mediated (p)ppGpp synthesis during heat and oxidative stress has been reported for diverse taxa, suggesting that the underlying mechanisms could be conserved [56–58]. However, little is known about the mechanism of Rel activation upon environmental stress.

RSH-type enzymes have been implicated in sensing and integrating many environmental cues beyond amino acid starvation. These additional signals may be transmitted by direct interaction of RSH-type with additional regulatory proteins, which expand and adapt SR-signalling to the respective requirements of the environmental niche [24, 25]. For example, growth inhibition in competent *B. subtilis* cells has recently been shown to be mediated by a specific interaction between Rel and ComGA, a membrane associated ATPase which is involved in uptake of DNA in competent cells [59]. It was suggested that the interaction of ComGA with Rel inhibits its hydrolase activity, resulting in the accumulation of (p)ppGpp and the inhibition of rRNA synthesis and growth. Furthermore, the growth-inhibition by the SR promotes increased antibiotic tolerance of competent cells and therefore contributes to bet-hedging and improves fitness of the population [59].

Our experiments demonstrate that Rel activation during heat- or oxidative stress can be inhibited by chloramphenicol similarly as during amino acid starvation (Fig. 1F, S1E). Therefore, the underlying activation mechanisms during environmental stress likely shares some similarities to the well-studied SR-activation upon amino acid deprivation and may also involve the sensing of uncharged tRNA on the ribosome [17,18,56,58].

Proteotoxic and oxidative stress results in the inactivation of labile enzymes and may thereby impair uptake or biosynthesis of certain amino acids and/or modulate the activity of aminoacyl-tRNA synthetases, resulting in an accumulation of uncharged tRNA which can serve as a signal to activate Rel [58, 60]. In addition, tRNAs and proteins of the translational machinery are prone to oxidation or modification upon stress, leading to translation stalling, which can also elicit the SR [61].

### The role of (p)ppGpp and the transcription factor Spx in global transcriptional regulation upon heat stress

Using RNA-seq, we observed large transcriptomic alterations mediated by (p)ppGpp in the Δ*rel* mutant (Fig. 3). Since *B. subtilis* lacks a DksA homolog, regulation of transcription by (p)ppGpp is achieved indirectly by lowering GTP levels, which reduces transcription of promoters that initiate with GTP. Accordingly, transcription of rRNA and r-protein genes was strongly reduced in the Δ*rel* strain as a consequence of the lower GTP level [28, 62]. In addition, the CodY activity is under allosteric control by cellular GTP levels [52]. While the effect of both mechanisms is prominently visible in Δ*rel* cells where the GTP level is strongly reduced, their impact is less noticeable in heat-stressed cells where a transient increase of alarmones, but no decrease of the GTP concentration, was observed (Fig. 1, S1). It is possible that the transient pulse, its kinetic and the generated total amount of (p)ppGpp induced by the raised temperature might not be sufficient to promote the strong reduction of cellular GTP that is observed during amino acid starvation (Fig. S1) [27]. It should be noted that a strong reduction of cellular GTP levels would most likely also interfere with the ability of *B. subtilis* cells to grow at 50 °C with a growth rate comparable to that at 37 °C (Fig. S9 B). When designing the RNA-seq experiment, we choose 48 °C as a simple heat shock condition for the mutant strains since it resembled the thermotolerance protocol (Fig. 1A) and the condition of previously published microarrays [14]. However, many phenotypes of Spx and (p)ppGpp could be observed best upon a stronger, but non-lethal, heat shock at 50 °C [14]. In contrast, while wildtype cells treated with 37/53 °C exhibit a strong increase of (p)ppGpp within the first minutes of stress (Fig 1D), the examination of cellular physiology is confounded by the rapid reduction of viability at this lethal condition (Fig. 2G) [5, 14].

We reported previously that transcription of rRNA can be down-regulated by the global regulator Spx. During heat stress however, down-regulation of rRNA was independent of Spx, suggesting that the loss of Spx was compensated by additional regulators [14]. Strikingly, we now observed that (p)ppGpp also engages in this down-regulation of rRNA during heat stress and that the concurrent activity of both Spx and (p)ppGpp is required to reach the full strength of this effect (Fig. 5A). This functional relationship of Spx and the SR is corroborated by the observation that the (p)ppGpp^0^ Δ*spx* mutant strain displays strongly impaired growth at both 37 °C and 50 °C (Fig. 5B). In addition, the observation that a (p)ppGpp^0^ *cxs*-2 strain, in which Spx can still up-regulate the stress response, exhibits a much less impaired growth than a (p)ppGpp^0^ Δ*spx* or (p)ppGpp^0^ *cxs*-1 strain in which Spx activity is fully disrupted. This suggests that at least either (p)ppGpp or Spx is required for the transcription of unknown factors necessary for efficient growth under adverse conditions. The observation that transcription of *spx* is also activated by (p)ppGpp via CodY in *Enterococcus faecalis* and *rel* transcription is activated by the disulfide-stress regulator σ^R^ in *Streptomyces coelicolor* points toward possible functional connection of these two regulators [63, 64].

Our RNA-seq dataset also indicates a possible activation of the SigB-dependent general stress response by (p)ppGpp during stress- and non-stress conditions (Fig S7). SigB becomes activated by decreased GTP levels as elicited by decoyinine [53, 54]. In addition, a requirement of L11, which is necessary for Rel synthetase activity, and Obg, a ribosome-associated GTPase that interacts with ppGpp, for the activation of SigB upon physical stress and an interaction of Obg with components of the SigB regulatory cascade was reported, suggesting an intricate connection between the ribosome, Rel and the general stress response [53,65,66].

### Control and protection of translation by (p)ppGpp and the role of Hpf during heat stress

Our results suggest that (p)ppGpp acts as a negative regulator of translation during heat shock. (p)ppGpp was shown to bind and inhibit many ribosome-associated GTPases thus interfering with ribosome assembly and arresting translation [32, 33]. The relative reduction of translation during stress would reduce the load on the protein quality control systems, thus alleviating the burden for cellular protein homeostasis upon protein folding stress [3, 67]. This hypothesis is supported by the observation that the (p)ppGpp^0^ mutant accumulated more protein aggregates during heat stress, whereas a Δ*rel* mutant strain exhibited significantly lower translation, but at the same time generated also significantly less protein aggregates (Fig. 2I). (p)ppGpp is also required for the efficient transcription and synthesis of the Hpf protein that promotes the formation of translationally inactive 100S disomes, which supports the fast regrowth of cells after stress conditions have ceased [43,68,69]. Furthermore, we observed that the 16S rRNA was degraded and translation was diminished in the (p)ppGpp^0^ mutant upon severe stress at 37/53 °C. This phenotype was neither rescued by *in trans* expression of *hpf* or the addition of antibiotics that inhibit translation, suggesting that a specific mechanism for the protective action of (p)ppGpp and not an inhibition of translation is required for this process. However, cells were reported to be able to fully recover from the heat-induced rRNA degradation [70]. It was recently observed *B. subtilis* that tRNA maturation defects could lead to an inhibition of rRNA processing and 30S assembly via the synthesis of (p)ppGpp [71]. These observations might be important to understand possible stress signaling pathways and also the protective effect of (p)ppGpp on translation under proteotoxic stress conditions.

### The role of the SR during the heat stress response

Taken together, our data suggest a model in which cells respond in a concerted manner to heat-mediated protein unfolding and aggregation, not only by raising the repair capacity, but also by decreasing translation to concurrently reduce the load on the cellular protein quality control systems (Fig. 7A). Upon heat shock, Rel is activated and rapidly synthesizes alarmones. These alarmones mediate, in conjunction with Spx, a strong down-regulation of ribosomal promoters together with the up-regulation of stress response-genes such as *hpf* during heat shock (Fig. 7B), while chaperones and stress-response genes controlled by Spx and other regulators are concurrently up-regulated. In addition, the second messenger (p)ppGpp could directly control the activity of translation factors and may thereby mediate a fast and immediate response to slow down translation during stress. Together, the combined readjustments on transcription and translation allow an efficient reallocation of cellular resources to the synthesis of stress response proteins and concurrently minimize the load on the protein quality control systems, thus contributing to protein homeostasis [3,67,72]. The unfolded protein response to misbalances in protein homeostasis in the endoplasmic reticulum of eukaryotic cells is a well-studied and analogous stress response mechanism where the up-regulation of chaperones is also coupled to the concurrent down-regulation of translation, albeit by different mechanisms [3, 73].

Interestingly, accumulation of (p)ppGpp upon heat or oxidative stress and its importance for stress resistance has also been reported in other *Firmicutes* and also *Proteobacteria* that differ widely in terms of (p)ppGpp signaling [56–58,74,75]. Accumulation of (p)ppGpp was shown to protect cells from salt or osmotic stress [76, 77]. Conversely, the lack of (p)ppGpp is known to renders cells sensitive to heat or oxidative stress [58,78,79], suggesting that activation of the SR, allowing the fast down regulation of translation, is an important and conserved part of the response to environmental stress in bacteria. It is interesting to note that SR was also implicated in *B. subtilis* competence development, facilitating a cellular state (also referred to as the K-state) where cells cease to divide, and most transcription and translation is strongly down-regulated. In these cells only competence proteins, together with DNA repair and recombination genes, are expressed, allowing the uptake and possible utilization of homologous of DNA in this specific cellular state of a subpopulation of stationary phase cells [59]. Bacterial cells thus appear to utilize the (p)ppGpp second messenger, which can interfere directly with basic cellular processes such as translation, replication and growth, as an important part of different regulatory networks, facilitating and allowing the survival of bacterial cells in fast changing environments with limited nutrient availability and exposure to various stress conditions.

## Methods

### Construction of strains and plasmids

Strains, plasmids and primers are listed in S1 Table. PCR-amplification and molecular cloning using *E. coli* DH5α as host was carried out according to standard protocols [80]. Point mutations were introduced via overlap-extension PCR. To generate pBSII-spxDD-spec, a fragment carrying *spx^DD^*, *lacI* and the spectinomycin resistance cassette was amplified from pSN56 [12] with primers p289/p223 and ligated using *Spe*I/*Nsi*I sites into the pBSIIE backbone amplified with primers p203/p288. Integrative plasmids were linearized by digestion with *Sca*I or *Bsa*I prior to transformation. Point mutations in the *rel* gene were first cloned in the pMAD vector and then re-amplified for cloning into pDR111.

Transformation of *B. subtilis* strains, the generation of scarless mutations using the pMAD system and the introduction of *cxs-1/2* mutations in *rpoA* was carried out as described previously [81–83]. Mutants were selected on 100 µg ml^−1^ spectinomycin, 10 µg ml^−1^ kanamycin, 1 µg ml^−1^ erythromycin, 25 µg ml^−1^ lincomycin or 5 µg ml^−1^ chloramphenicol, respectively. To obtain the (p)ppGpp^0^ strain (BHS214), markerless *sasA*^E154V^ and *sasB*^E139V^ mutations were introduced into *B. subtilis 168* cells by successive transformation and recombination of plasmids pMAD-sasA^E154V^ and pMAD-sasB^E139V^, yielding strain BHS204. Next, a PCR amplified fragment carrying *rel::erm* [22] and flanking homologous regions was transformed to generate BHS214. Since the (p)ppGpp^0^ strain fails to develop natural competence, additional mutations were introduced in BHS204 and transformed with a PCR-amplified *rel::erm* fragment or BHS214 genomic DNA in a second step.

### Growth conditions

*B. subtilis* strains were grown in LB medium (5 g L^−1^ yeast extract, 10 g L^−1^ tryptone-peptone, 10 g L^−1^ NaCl) or minimal medium [84] supplemented with 0.5 % casamino acids in water baths with 200 rpm orbital shaking at the desired temperatures. 1 mM IPTG or 0.4 % xylose was supplemented if required.

### Survival and viability assays

The assays for thermotolerance development, survival and preparation of protein aggregate are described previously [5]. 1 mM IPTG was added to induce expression of recombinant proteins 30 min before the division of the culture. The influence of decoyinine on thermotolerance was tested in 1.5 mL tubes in thermoshakers. Detection of aggregates by fluorescence microscopy was described previously in [41]. Spot colony formation assays were carried out as described previously and incubated at the indicated temperatures [14].

### Transcription analysis

Strains were grown in LB and treated as indicated. Samples of 15-25 mL were harvested by centrifugation for 3 min at 3.860 xg at 4 °C and frozen in liquid nitrogen. Isolation of total RNA, treatment with DNase I (NEB) and quality control by native agarose gel electrophoresis, methylene blue staining and northern blotting was described previously [14]. Northern blotting, hybridization with DIG-labeled RNA probes and detection was carried out as described previously [14]. Primers for the synthesis of probes are listed in S1 Table. Reverse transcription and qPCR were carried out as described previously [14]. The primers are listed in S1 Table. 23S rRNA was used as a reference.

### RNA sequencing

Cells of BHS220, BHS319 and BHS368 were grown in 150 mL LB medium in 500 mL flasks in water baths at 37 °C and 200 rpm. In the mid-exponential phase (OD_600 nm_ ∼ 0.4), the culture was divided and shifted to 48 °C or left at 37 °C. After 15 min, samples were withdrawn and both cultures were shifted to 53 °C for another 15 min and harvested. Cells from 25 mL medium were pelleted by centrifugation for 3 min at 3.860 × g and 4 °C and flash-frozen in liquid nitrogen. RNA was prepared the using phenol/trizol method as described in [85] and treated with TURBO DNase (Invitrogen). RNA quality was assessed on a Bioanalyzer 2100 System (Agilent).

rRNA depletion from total RNA using MICROBExpress (Ambion), treatment with tobacco acid pyrophosphatase (TAP) for +TAP libraries, library preparation, Illumina sequencing and quality control of the sequencing output was carried out as described previously [86]. Reads were mapped to the *Bacillus subtils* 168 genome with insertion of *rrnJp1-lacZ* in the *amyE* site (strain BHS220, *amyE::rrnJp1-lacZ cat*) using Bowtie2 (version 2.1.0) reads [87] with default parameters and filtered for uniquely mapped reads using SAMtools [88]. The DEseq2 package with default parameters was used for the detection of differentially expressed genes from raw count data of triplicate experiments [89]. Expression changes were considered significant if differentially regulated by at least 4-fold (*p*-value ≤ 0.05). The data have been deposited in NCBI’s Gene Expression Omnibus and are accessible through GEO Series accession number GSE125467 [90]. Identification of transcription start sites (TSS) and gene set enrichment analysis (GSEA) is described in S1 Text.

### *In vitro* transcription

*In vitro* transcription assays using purified *B. subtilis* RNA polymerase and Spx protein was carried out as described previously [14].

### Fluorescence microscopy

Strain BIH369 (*lacA::Pxyl-yocM-mCherry erm*) was grown in LB medium + 0.5 % xylose. The culture was divided in the mid-exponential phase, supplemented with puromycin for 15 min and subjected to fluorescence microscopy in a Axio Imager.Z2 (Zeiss) microscope using the RFP filter set [14].

### SDS PAGE and Western blotting

Strains were grown in LB medium and treated as indicated, harvested by centrifugation for 5 min at 3.860 xg at 4 °C, washed in TE buffer (10 mM TRIS-HCl, 1 mM EDTA, pH 8.0) and disrupted by sonication in TE supplemented with 1 mM PMSF. Equal amounts of protein were separated by SDS-PAGE and stained with coomassie or subjected to western blotting [91–93]. For signal detection, polyclonal α-Hpf antibody (1:5000) [69] or monoclonal anti-puromycin antibody (1:10.000, Merck) and HRP-conjugated anti-mouse or anti-rabbit antibodies (1:10.000, Roth) were used in conjunction with the ECL-system as described previously [14]. Images were acquired using a ChemoStar Imaging System (Intas, Göttingen, Germany)

### Translation rate analysis

Strains were grown in LB medium and treated as indicated. For *in vivo* labeling, 10 mL medium were separated, supplemented with 1 µg mL^−1^ puromycin (Roth) and incubated for 15 min at the same conditions. Then, samples were supplemented with 25 µg mL^−1^ chloramphenicol, harvested by centrifugation for 5 min at 3.860 xg at 4 °C, washed in TE buffer (10 mM TRIS-HCl, 1 mM EDTA, pH 8.0) and disrupted by sonication in TE supplemented with 1 mM PMSF. Equal amounts of protein were directly spotted on nitrocellulose membranes (5 µg) or subjected to SDS-PAGE and western blotting [80]. Puromycin-signals were detected using monoclonal anti-puromycin antibody (1:10.000, Merck), HRP-conjugated anti-mouse antibody (1:10.000, Roth) and the ECL-system in a ChemoStar imaging system (Intas, Göttingen, Germany). Signals were analyzed using Fiji distribution of ImageJ [94].

### Sucrose density gradient centrifugation analysis

Early exponential phase cultures of *B. subtilis* strains grown in LB medium were treated with heat shock at 48 °C or 48 °C/53 °C for 15 min each. Samples of 50 mL were supplemented with 50 µg mL^−1^ chloramphenicol to stall translation and harvested by centrifugation at 4000 × g for 10 min at 4 °C. Cells were resuspended in 25 mM HEPES-KOH, pH 7.5, 150 mM KOAc, 25 mM Mg(OAc)_2_, 1 mM dithiothreitol (DTT), n-Decyl−β−D-thiomaltopyranoside (DTM), 5 % (w/v) sucrose) and lysed by sonication. The lysate was cleared by centrifugation at 16,000 × g for 15 min at 4 °C. 10 OD_260_ units were loaded on a 10 mL 5-45 % (w/v) sucrose gradient prepared in the same buffer, run in a SW-40 Ti rotor (Beckman Coulter) at 57471 × g for 16.5 h and analyzed using a Gradient Station (Biocomp) with an Econo UV Monitor (Bio-Rad).

### Quantification of nucleotides

Cells were grown in minimal medium supplemented with 0.5 % casamino acids to support the growth of (p)ppGpp deficient strains [95] and treated as indicated. Samples of 2 mL were removed, supplemented with 75 µL 100 % formic acid and incubated on ice for 30 min. Extraction of nucleotides was carried out as described in [96] and detected by HPLC-ESI-MS/MS on a QTRAP 5500 instrument. Analytes were separated on a Hypercarb column (30 × 4.6 mm, 5 µm particle size) in a linear gradient of solvent A (10 mM ammonium acetate pH 10) and solvent B (acetonitrile) at a flow rate of 0.6 mL/min from 96 % A + 4 % B (0 min) to 40 % A + 60 % B (8 min) into the ESI ion source at 4.5 kV in positive ion mode. Tenofovir was used as internal standard. pGpp and pppGpp standards were synthesized *in vitro* from ATP and GTP or GMP as described previously [97]. ppGpp was purchased from Trilink Biotechnologies.

## Acknowledgements

We thank Dr. Laurent Jannière (<u>French National Centre for Scientific Research</u>, Paris, France) and the National BioResource Project (National Institute of Genetics, Japan) for providing strains and plasmids.

## Supporting information captions

**S1 Fig.: Alarmone and GTP levels during stress and starvation.**

**(A)** Means and SEM of GTP after the application of different stress conditions. Sample sizes and treatments are the same as in Fig. 1 A, B. NV: DL-norvaline, SHX: serine hydroxamate. Asterisks (*) indicate significance (*p_adj._* ≤ 0.05) of combined pGpp, ppGpp and pppGpp levels according to the Kruskal-Wallis and Dunn-Bonferrroni test. **(B)** Levels of GTP during thermotolerance development. Wildtype cells were grown at 37 °C and shifted to 48 °C for 15 min (pre-shock), then to 53 °C or directly to 53 °C. Samples were taken at 2, 5 and 15 min. Means and SEM of four independent experiments are shown. All changes are not significant (p ≤ 0.05) according to the Kruskal-Wallis test. **(C)** Means and SEM of GTP levels during heat shock in of wildtype cells or strains with mutations in (p)ppGpp synthetases (*sasA/B*^−^: BHS204, *rel*^E324V^: BHS709; (p)ppGpp^0^: BHS214). Sample sizes are the same as in Fig. 1E. Asterisks indicate significant changes (*p* ≤ 0.05) according to Welch’s *t-*test. **(D)** Relative changes in the transcription of *hpf* during heat shock in the same strains (15 min 50 °C). Means and SEM of three independent experiments are shown. Asterisks (*) indicate significance (*p_adj._* ≤ 0.05) according to the Kruskal-Wallis and Dunn-Bonferrroni test. **(E)** The influence of chloramphenicol on GTP levels during stress. Sample sizes and treatments are the same as in Fig. 1F. Asterisks indicate significant changes (*p* ≤ 0.05) according to Welch’s *t-*test.

**S2 Fig.: Phenotype of single deletions of (p)ppGpp synthetase genes.**

**(A)** Cellular GTP levels in wildtype or Δ*rel*: (BHS126) strains. **(B)** Growth of strains with mutations or deletions in (pp)pGpp metabolizing enzymes in rich LB medium. Δ*rel*: BHS126, (p)ppGpp^0^: BHS214. **(C/D)** Survival of wildtype (black lines) and mutant strains (Δ*sasB*: BHS127 or Δ*sasA*: BHS128) red lines at 53 °C with (48/53 °C) or without (37/53 °C) pre-shock. Means and SEM of at least three independent experiments are shown. Open symbols: no pre-shock, closed symbols: 15 min pre-shock at 48 °C. **(E)** Growth of wildtype cells or strains with deletions in *sasB* (BHS127) or *sasA* (BHS128) on agar plates at 37 °C, during heat stress (55 °C) or oxidative stress (0.2 mM diamide).

**S3 Fig.: Thermotolerance and survival of strains expressing *rel* variants *in trans*.**

**(A)** Levels of alarmones in these strains after the application of 1 mM IPTG for 15 min. Asterisks indicate significant changes (*p* ≤ 0.05) of combined alarmone levels according to Welch’s *t-*test. **(B-E)** Survival of wildtype (black lines) and mutant strains (red lines) at 53 °C without pre-shock (37/53 °C; open symbols) or with pre-shock (15 min 48 °C/53 °C; closed symbols). Means and SEM of at least three independent experiments are shown. Strains were supplemented with 1 mM IPTG 15 min prior to temperature shift. **(B)** Expression of a truncated, hyperactive *rel* variant from *E. coli* (designated *relA_hyper_*). **(C)** Expression of *rel_B.s._* with inactive hydrolase domain (E77A D78A) in the (pp)pGpp^0^ strain. **(D)** Expression of a truncated, inactive *relA* variant from *E. coli* (*relA_inactive_*). **(E)** Expression of *rel_B.s._* with inactive synthetase domain (E324V) in the (pp)pGpp^0^ strain.

**S4 Fig.: thermotolerance and survival of strains expressing treated with decoyinine** Thermotolerance development and survival of wildtype cells treated with decoyinine (red lines) or left untreated (black lines). Means and SEM of at least three independent experiments are shown. Strains were supplemented with 50, 250, 400 or 1000 µg ml^−1^ decoyinine 15 min before heat treatment. Open symbols: no pre-shock, closed symbols: 15 min pre-shock at 48 °C. n.d.: not determined, no cfu could be detected from 100 µl cell culture.

**S5 Fig.: Analysis of transcription start sites**

**(A)** Venn Diagram showing the overlap of TSS identified in this study with transcription upshifts identified in [98]. **(B)** Venn Diagram depicting the classification of the identified TSS. **(C)** Length distribution of the distance from the TSS to the translation initiation site. **(D)** Sequence logos of the region around the TSS of genes up- or down-regulated during different conditions. **(E)** Predicted sigma factors in the different TSS classes.

**S6 Fig.: Global differences in gene expression of heat shocked**

The distributions of all up- and down-regulated genes for the indicated conditions are shown. Bar tracks indicate the distribution of the respective functional groups. **(A)** Wildtype cells (BHS220) heat shocked at 48 °C or 53 °C versus unstressed cells. **(B)** Wildtype (BHS220) versus (p)ppGpp^0^ cells (BHS319) at37 or 48 °C. **(C)** wildtype (BHS220) versus Δ*rel* cells (BHS368) at 37 °C or 48 °C.

**S7 Fig.: Up- or down-regulation of regulons or gene categories.**

Points in the scatterplot represent log2-transformed up- or down-regulation of individual genes of the respective regulons relative to wildtype cells at 37 °C. Blue/gray color indicates transcriptional changes above/below the significance threshold (see Materials and Methods). Horizontal bars represent the median expression changes of the whole gene set.

**S8 Fig.: (p)ppGpp mediated transcriptional changes during heat stress**

Relative changes in the transcription of selected genes during heat shock at 50 °C in wildtype (BHS220), (p)ppGpp^0^ (BHS319) and Δ*rel* (BHS368) strains determined by RT-qPCR. Means and SEM of three replicates are shown. Asterisks indicate significance (*p* ≤ 0.05) according to Welch’s *t-*test, *n.s.*: not significant.

**S9 Fig.: (pp)pGpp and Spx act complementary.**

**(A)** RT-qPCR experiment showing the relative transcription of *rplC* and *rplO* in wildtype (BHS220), Δ*spx* (BHS222), (pp)pGpp^0^ (BHS319) or (pp)pGpp^0^ Δ*spx* (BHS766) cells treated with or heat stress at 50 °C for 15 min. Means and SEM of three replicates are shown. Asterisks indicate significant changes (*p* ≤ 0.05) of transcript levels according to Welch’s *t-* test, **(B)** Growth of wildtype (BHS220), Δ*spx* (BHS222), (pp)pGpp^0^ (BHS319), (pp)pGpp^0^ Δ*spx* (BHS766), (pp)pGpp^0^ cx*s*-1 (BHS954) or (pp)pGpp^0^ cx*s*-2 (BHS949) cell in LB medium at 37 °C or 50 °C. **(C)** The fraction of aggregated proteins (left) or soluble proteins (right) in wildtype, Δ*spx* (BHS014), (pp)pGpp^0^ (BHS214) or (pp)pGpp^0^ Δ*spx* (BHS766) cells treated with or heat stress at 50 °C for 15 min. **(D)** Relative translation of a strain carrying an inducible copy of Spx^DD^ (BHS201) with and without the addition of IPTG. Means and SEM of seven independent experiments are shown. Asterisks indicate significance (*p* ≤ 0.05) according to Welch’s *t-*test.

**S10 Fig.: (pp)pGpp and Spx act independently.**

**(A)** Northern and western blot of wildtype, Δ*spx* (BHS014) or (pp)pGpp^0^ (BHS214) strains treated with or without DL-norvaline. Cells were grown in minimal medium supplemented with 0.5 % casamino acids to OD_600_ 0.4. The medium was removed by centrifugation and the cells were resuspended in fresh medium with casamino acids (--) or 0.5 mg/ml DL-norvaline (+) and grown for 30 min. **(B)** Relative transcription of *rrnJp1-lacZ* with or without expression of *spx^DD^* with 1 mM IPTG for 30 min in the wildtype or (pp)pGpp^0^ background. Means and SEM of three replicates are shown. **(C)** *in vitro* transcription from selected promoters with or without Spx and ppGpp under reducing (+ DTT) or oxidizing (- DTT) conditions. Means and SEM of three replicates and a representative autoradiogram are shown.

**S11 Fig.: Puromyin labels nascent proteins and does not disturb protein homeostasis at low concentration.**

**(A)** Accumulation of subcellular protein aggregates (fluorecent spots) after the addition of puromycin visualized by YocM-mCherry. BIH369 cells were grown in LB + 0.5 % xylose and treated with 1, 10 or 25 µg ml^−1^ puromycin or left untreated for 15 min. Phase contrast images (P.C.) and fluorescence images with RFP-filters (YocM-mCherry) are shown. **(B)** The effect of puromycin on growth. Wildtype cells were grown in LB to the mid-exponential phase (OD_600_ 0.4) and supplemented with puromycin at the indicated concentrations. **(C)** Dot blot or western blot of puromycin-labeled proteins. Exponentially growing cells grown in LB were treated with the indicated concentrations of puromycin for 15 min. **(D)** Outline of the genotypes of the RIK1066 strain, carrying an inducible copy of *sasA* in the (p)ppGpp^0^ background. **(E)** Relative puromycin incorporation in RIK1066 cells treated with or without 1 mM IPTG. Cells were incubated with 1 mg ml^−1^ puromycin for 15 min, added directly to the medium after the addition of IPTG (0-15 min) or after 15 min (15-30 min), then harvested. One representative experiment and means and SEM from the quantification of three independent experiments are shown. Asterisks indicate significance (*p* ≤ 0.05) according to Welch’s *t-*test.

**S12 Fig.: Relative translation of heat shocked cells.**

**(A)** Relative translation (puromycin incporpration) of wildtype, (p)ppGpp^0^ (BHS214) and Δ*rel* (BHS126) strains during heat stress at 53 °C. 1 µg ml^−1^ puromycin was added for 15 min to the medium directly after (0-15 min) or 15 min after the temperature upshift. Means and SEM of three independent experiments are shown. Asterisks indicate significance (*p* ≤ 0.05) according to Welch’s *t-*test. **(B)** Representative experiment from Fig. 6 B & S9 A. **(C)** Methylene blue stained membranes showing the integrity or degradation of rRNA after severe heat stress (53 °C). Wildtype, (p)ppGpp^0^ (BHS214) or Δ*rel* (BHS126) cells were heat-shocked at 48 °C, 53 °C or 48/53 °C for 15 min each. 2 µg total RNA was separated on denaturing agarose gels and blotted on nylon membranes. **(D)** Western blot showing Hpf levels during thermotolerance development in wildtype, (p)ppGpp^0^ (BHS214) or Δ*rel* (BHS126) strains. Cells were heat shocked for 15 min each at the indicated temperature(s). **(E)** Methylene blue stained membranes showing the integrity or degradation of rRNA. Wildtype, (p)ppGpp^0^ (BHS214) Δ*hpf* (BHS008) or (p)ppGpp^0^ P_spac_-*hpf* (BHS626) cells were treated with or without heat shock at 53 °C for 15 min. 1 mM IPTG was added to the strains to induce the expression of *hpf* 15 min prior to heat shock. 2 µg total RNA was separated on denaturing agarose gels and blotted on nylon membranes. **(F)** Wildtype, (p)ppGpp^0^ (BHS214) (p)ppGpp^0^ P_spac_-*rel* (BHS622) or (p)ppGpp^0^ P_spac_-*hpf* (BHS626) were spotted on agar plates supplemented with 1 mM IPTG and incubated over night at 37 °C or 55 °C. **(G)** rRNA degradation after severe heat stress (53 °C) in wildtype or (p)ppGpp^0^ (BHS214) cells left untreated or treated with 5 µg ml^−1^ chloramphenicol or 100 µg ml^−1^ spectinomycin 15 min prior to the application of stress. 2 µg total RNA was separated on denaturing agarose gels and blotted on nylon membranes.

**S1 Table: List of strains, plasmids and oligonucleotides.**

This table lists all *B. subtilis* strains, plasmids and oligonucleotides used in this study.

**S1 Text: List of identified transcription start sites.**

In this document, additional methods and results are described.

**S1 Dataset: List of identified transcription start sites.**

In this dataset, all identified transcriptional start sites and their classification is shown.

**S2 Dataset: Results of the gene set enrichment analysis.**

This dataset lists all enriched functional categories and regulons for each for each condition in separate sheets.

**S3 Dataset: List of differentially expressed genes**

Global gene expression changes for all conditions are listed in separate sheets.

